# Alterations in Ion Channel Expression Surrounding Implanted Microelectrode Arrays in the Brain

**DOI:** 10.1101/518811

**Authors:** Joseph W. Salatino, Arya P. Kale, Erin K. Purcell

**Affiliations:** Department of Biomedical Engineering, Michigan State University; Institute for Quantitative Health Science & Engineering (IQ), Michigan State University; Department of Electrical and Computer Engineering, Michigan State University; Neuroscience Program, Michigan State University

**Keywords:** Neuroprostheses, tissue response, brain-machine interfaces, ion channels, plasticity, microelectrode arrays

## Abstract

Microelectrode arrays designed to map and modulate neuronal circuitry have enabled greater understanding and treatment of neurological injury and disease. Reliable detection of neuronal activity over time is critical for the successful application of chronic recording devices. Here, we assess device-related plasticity by exploring local changes in ion channel expression and their relationship to device performance over time. We investigated four voltage-gated ion channels (Kv1.1, Kv4.3, Kv7.2, and Nav1.6) based on their roles in regulating action potential generation, firing patterns, and synaptic efficacy. We found that a progressive increase in potassium channel expression and reduction in sodium channel expression accompanies signal loss over 6 weeks (both LFP amplitude and number of units). This motivated further investigation into a mechanistic role of ion channel expression in recorded signal instability. We employed siRNA in neuronal culture to find that Kv7.2 knockdown (as a model for the transient downregulation observed at 1 day in vivo) mimics excitatory synaptic remodeling around devices. This work provides new insight into the mechanisms underlying signal loss over time.

## Introduction

Charge movement across the cell membrane through ion channels enables the conduction and propagation of electrical signals that underlie neuronal communication and function^1^. The remarkable diversity of ion channels in the mammalian brain (comprising more than 90 voltage-gated potassium channels alone) facilitates the rich repertoire of excitable properties that shape neuronal signaling to encode information along neuronal networks^1,2^. The effective use of microelectrode arrays implanted in the brain relies on the ability to record electrical signals from single neurons and their populations over time^3–6^. Neuronal loss and glial encapsulation are well-known consequences of implanting commonly used electrode designs^7–9^, but impacts on the residual function of remaining neurons are unknown. Ion channel expression and function is highly dynamic and modulated by many factors^1^, including changes to the surrounding environment caused by injury^10–13^ and inflammation^14–16^. Channel modulation can impact not only the signal generation capabilities of single neurons, but also their frequencies, patterns, and waveform characteristics that underlie information encoding^1,2^. Channel modulation can also contribute to neuronal network dysfunction (e.g., transcriptional and post-translational channel effects of cytokine exposure can result in cortical circuit hyperexcitability and epileptogenesis^17^). Therefore, injury caused by device insertion could influence the signal detection of microelectrodes by modifying the firing properties and coordinated function of surrounding neurons over time.

Several lines of evidence support the notion that injury and inflammation associated with device insertion could result in changes to the structure and function of nearby neurons. Cytokines and gliotransmitters released by reactive astrocytes have been shown to impact neuronal health (neurotoxic/protective effects^18–21^) and function (ion channel/synaptic remodeling^17,22–24^) to modify the composition, connectivity and excitability of local neuronal networks^17,18,23,24^. Inflammatory cytokines possess neuromodulatory properties that alter ion channel expression and function in neuronal circuits that develops over acute and chronic periods of time^16,17^. Although results vary^25^, general observations follow a trend from acute hyperexcitability to chronic hypoexcitability within affected neuronal networks^17^. Similar trends have frequently been observed following traumatic brain and axonal injury models, where shifts in excitation/inhibition likewise occur^26–28^. Interestingly, axonal damage produces transient changes in electrophysiological properties of both axotomized and surrounding intact neurons in the injured cortex^29^, where transient increases in membrane potentials (~10mV) occurred within initial days that are of sufficient magnitude to impact the signal detection capabilities of implanted electrode arrays^30^ (where a ~10mV intracellular amplitude difference can equate to ~70uV extracellular amplitude difference^30^). The authors attributed these effects to changes in ion channel expression and function in axonal compartments (specifically, sodium channel and A-type potassium channel expression^29^).

In this work, we have developed a platform for assessing local changes in ion channel expression surrounding implanted functional electrode arrays over time. While recognizing that neurons express a diverse repertoire of ion channels, we have chosen to initially explore four voltage-gated ion channels (Kv1.1, Kv4.3, Kv7.2, and Nav1.6) based on their roles in regulating action potential generation^31^, firing patterns^32–34^, and synaptic efficacy^33^ (**Table 1**). Nav1.6 has been implicated in electrophysiological abnormalities following axonal injury^29^, where induced channel alterations have been demonstrated following axonal trauma^35^, traumatic brain injury^10^, and exposure to inflammatory cytokines^36^ that can evolve over time^17,37,38^. Likewise, A-type potassium channels (e.g., Kv4.3/Kv4.2) have been proposed to contribute to the loss of intrinsic bursting activity surrounding axotomized neurons^29^, where expression is transiently downregulated following traumatic brain injury^12^. Upregulated Kv1.1 expression at 6-8 weeks following CNS injury^39^ has been shown to be a mechanism for axonal dysfunction in surviving axons, where increased K+ conductance was proposed to act as a shunt for blocking axonal conduction^39,40^. Finally, Kv7.2 regulates vesicular glutamate transporter 1 (VGLUT1) expression and acts as a brake for repetitive firing^33^, where our group observed changes in VGLUT1 expression surrounding implanted microelectrodes over time that motivated further investigation into a mechanistic role of this channel^41^. Here, we report a progressive elevation in potassium channel expression coupled with a loss of sodium channel expression surrounding devices. These changes accompany a loss of signal over 6 weeks. Further, we provide insights into a mechanistic role of these ion channels in signal loss using siRNA in culture. Our study shows novel mechanisms of plasticity surrounding implanted devices that may affect their signal instability and long-term performance.

**Table 1.** Motivation for ion channel selection.

| **Ion Channel** | **Channel Type** | **Functional Role** | **Motivator** |
| --- | --- | --- | --- |
| Nav1.6 | Most abundant Na+ channel / clustered at axon hillock | Initiating action potentials (depolarization) | Down-regulation shown to induce hypoexcitability<sup>42</sup> |
| Kv1.1 | Delayed rectifier | Setting action potential threshold / for AP down-stroke | Blocking/KO shown to induce hyperexcitability<sup>32</sup> |
| Kv4.3 | A-type / inactivating | Setting inter-spike interval/firing rate | Blocking/KO shown to induce hyperexcitability<sup>43</sup> |
| Kv7.2 | M-type | Regulating synaptic transmission / acts as a brake for repetitive firing | ↓I<sub>m</sub>(M-current)<br>↑excitatory synaptic density<sup>33</sup> |

## Results

### Ion channel expression evolves over time

Based on motivations described in **Table 1**, we chose to explore whether shifts in the expression of selected ion channels occurs at the interface of implanted single-shank microelectrode arrays over 6 weeks using quantitative immunohistochemistry (with time points at 1 day, 1 week and 6 weeks). Images obtained using confocal laser scanning microscopy (**Fig. 1**) were analyzed using a custom-modified MATLAB script as previously reported^41^. Briefly, ion channel expression intensity was analyzed as a function of distance from the insertion site, where fluorescence intensity was calculated within 10um bins that were generated to extend radially from the user-defined insertion site (a total of 27 bins spanning a 270um radius). The same secondary antibody was used for all ion channels and distinct spatiotemporal patterns of expression were observed for each channel, mitigating the likelihood that non-specific background labeling contributed to our results.

**Figure 1.**
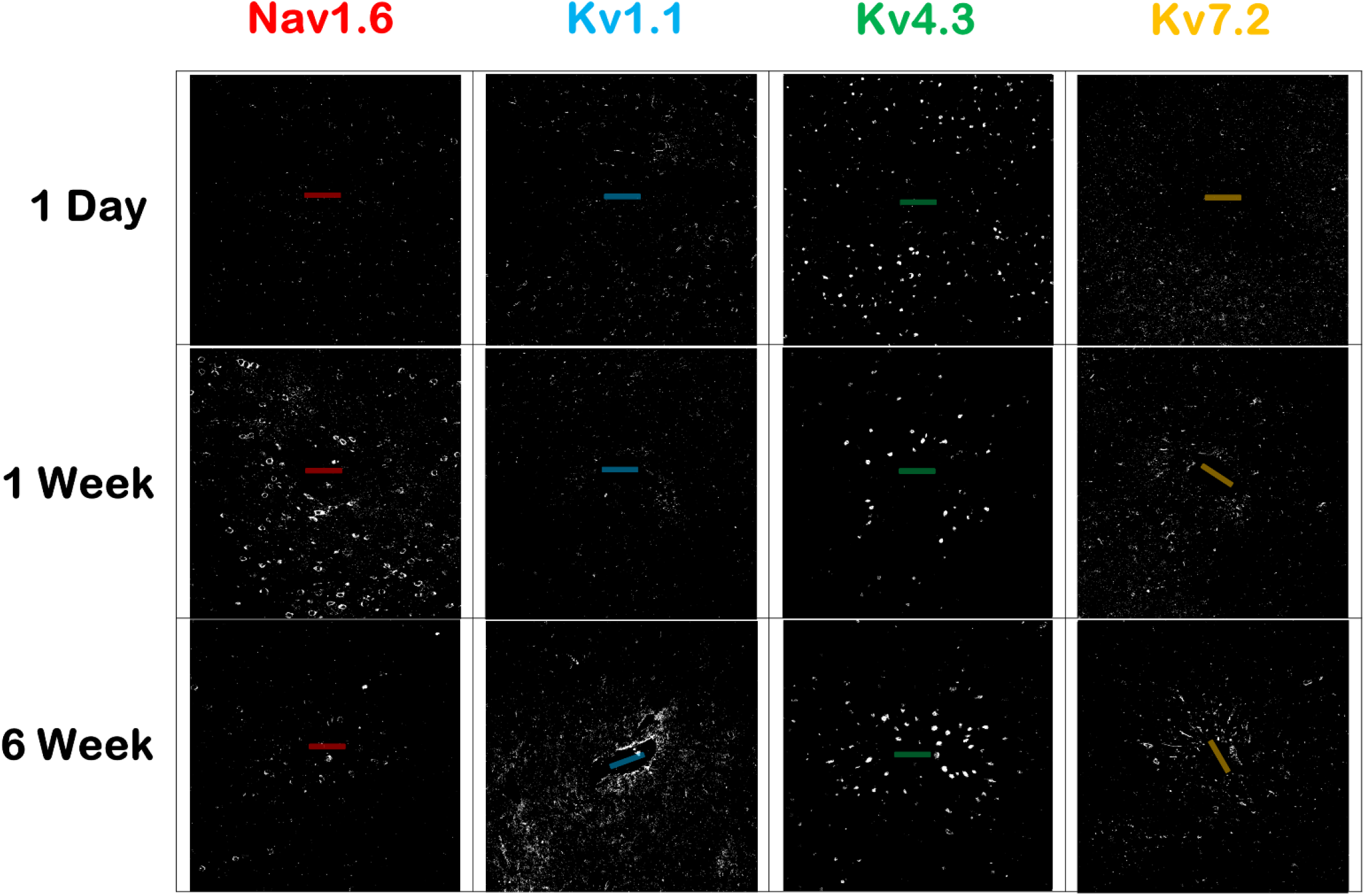
Confocal laser scanning microscopy of ion channel expression surrounding the insertion site. Example images of ion channel expression surrounding the device tract. Immunohistochemistry reveals fluorescently stained ion channels on horizontal tissue sections taken from layer V of the primary motor cortex using the same secondary antibody. Electrodes illustrated for reference with dimensions to scale (100um x 15um).

#### Spatial differences in expression

To assess spatial differences between stains at each time point, we normalized intensity bins for each stain to their respective final bins as previously reported^41^ and began with comparing the first 40um to the last 40um for statistical significance using a linear mixed effects model (**Fig. 2A**). At 1 day, we observed a significant reduction in both Kv7.2 (*) and Kv4.3 (***), followed by significant elevations in Kv7.2, Kv4.3, and Nav1.6 (***) at 1 week, and finally significant elevations in Kv7.2, Kv4.3, and Kv1.1 (***) at 6 weeks. Early local reductions in potassium channel expression at 1 day are followed by robust elevations at 1 and 6 weeks, and an elevation in Nav1.6 expression at 1 week is subsequently reduced by 6 weeks. The results reveal a progressive increase in potassium channel expression coupled with a reduction in sodium channel expression surrounding devices over 6 weeks.

**Figure 2.**
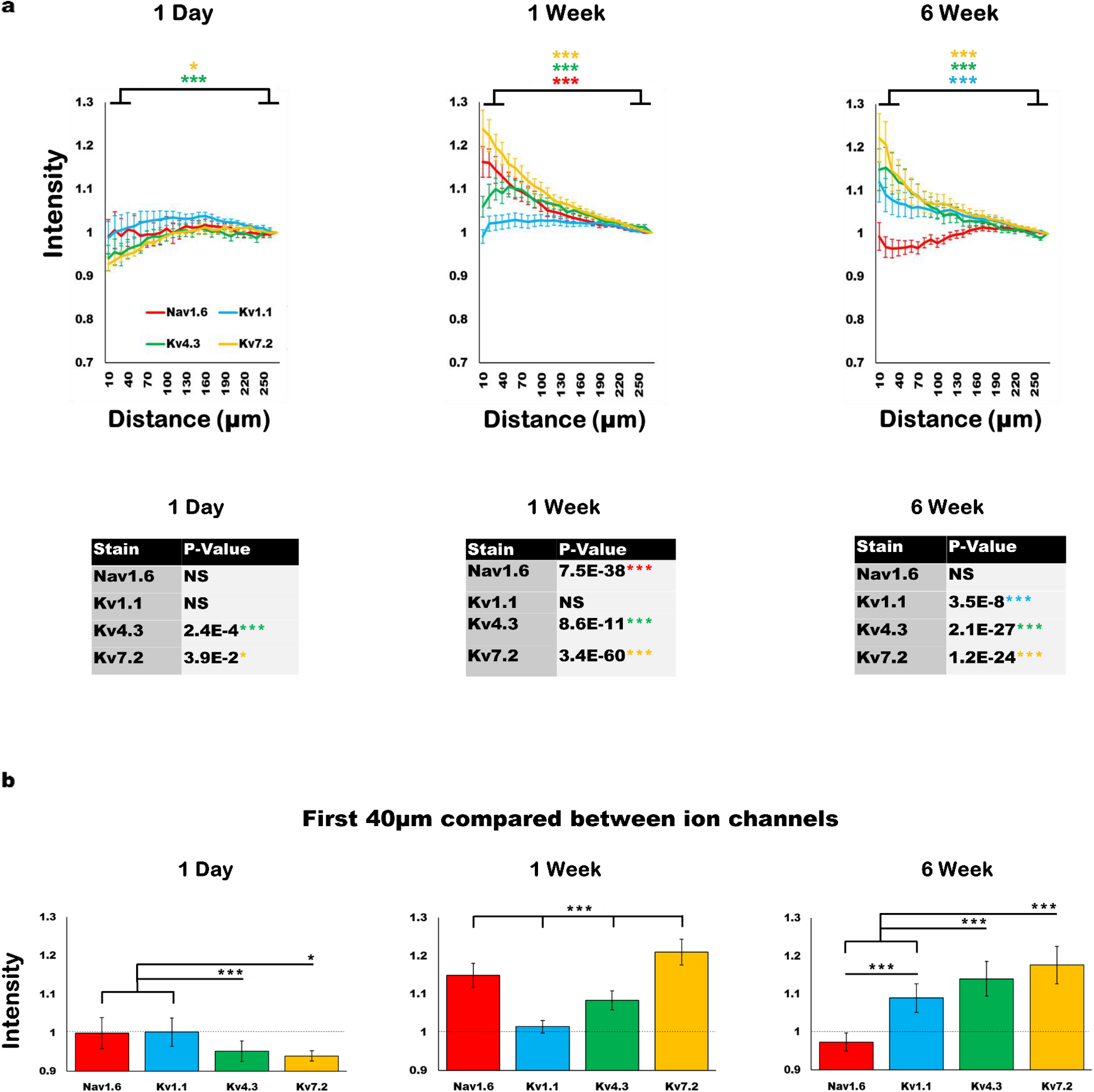
Spatial differences in expression at each time point: Progressive increase in potassium channel expression is coupled with a reduction in sodium channel expression over 6 weeks. **A)** Averaged intensity from ion channel expression (normalized to final bin) revealed an increase in potassium channel expression and a loss of sodium channel expression over 6 weeks (p-values comparing 0-40um and 230-270um depicted). **B)** Significance compared between 0-40um of each ion channel. Significance depicted as *p<0.05 and ***p<0.001. “NS” denotes non-significance. Standard error bars depicted in both panels. For each ion channel, there was an average of 7 devices and 21 tissue sections analyzed per time point.

Next, we compared the first 40ums between channels at each time point for statistical significance, as depicted in **Fig. 2B**. Although represented with bar graphs for visual ease, these results still incorporated distance-related effects using the same mixed model in Fig. 2A (each bar represents the averaged value for the first 40um of the given stain). At 1 day, both Nav1.6 and Kv1.1 were statistically different from Kv7.2 (*) and Kv4.3 (***), followed by significant differences between all ion channels at 1 week (***). At 6 weeks, Nav1.6 was significantly different from all other ion channels (***) and Kv1.1 was significantly different from both Kv7.2 and Kv4.3 (***) (**Fig. 2B**). The results support a shift toward a decrease in sodium channel expression and an increase in potassium channel expression over the chronic 6-week time course.

#### Temporal differences in expression

To investigate temporal differences in expression levels, we normalized 1 and 6 week expression values to 1 day expression values (bin-for-bin) and displayed the results as a relative percentage change (**Fig. 3**). To quantify temporal shifts, we calculated the area under the curve to assess the relative percentage change for the total area for each ion channel (**Fig. 3B**). At 1 week, the total integrated area revealed a relative decrease in Nav1.6 (−12%), and a relative increase in Kv1.1, Kv4.3 and Kv7.2 (94%, 175%, and 255%, respectively). At 6 weeks, the total area showed a greater relative decrease in Nav1.6 (−154%), and a sustained relative increase in Kv1.1, Kv4.3 and Kv7.2 channels (98%, 97% and 180%, respectively).

**Figure 3.**
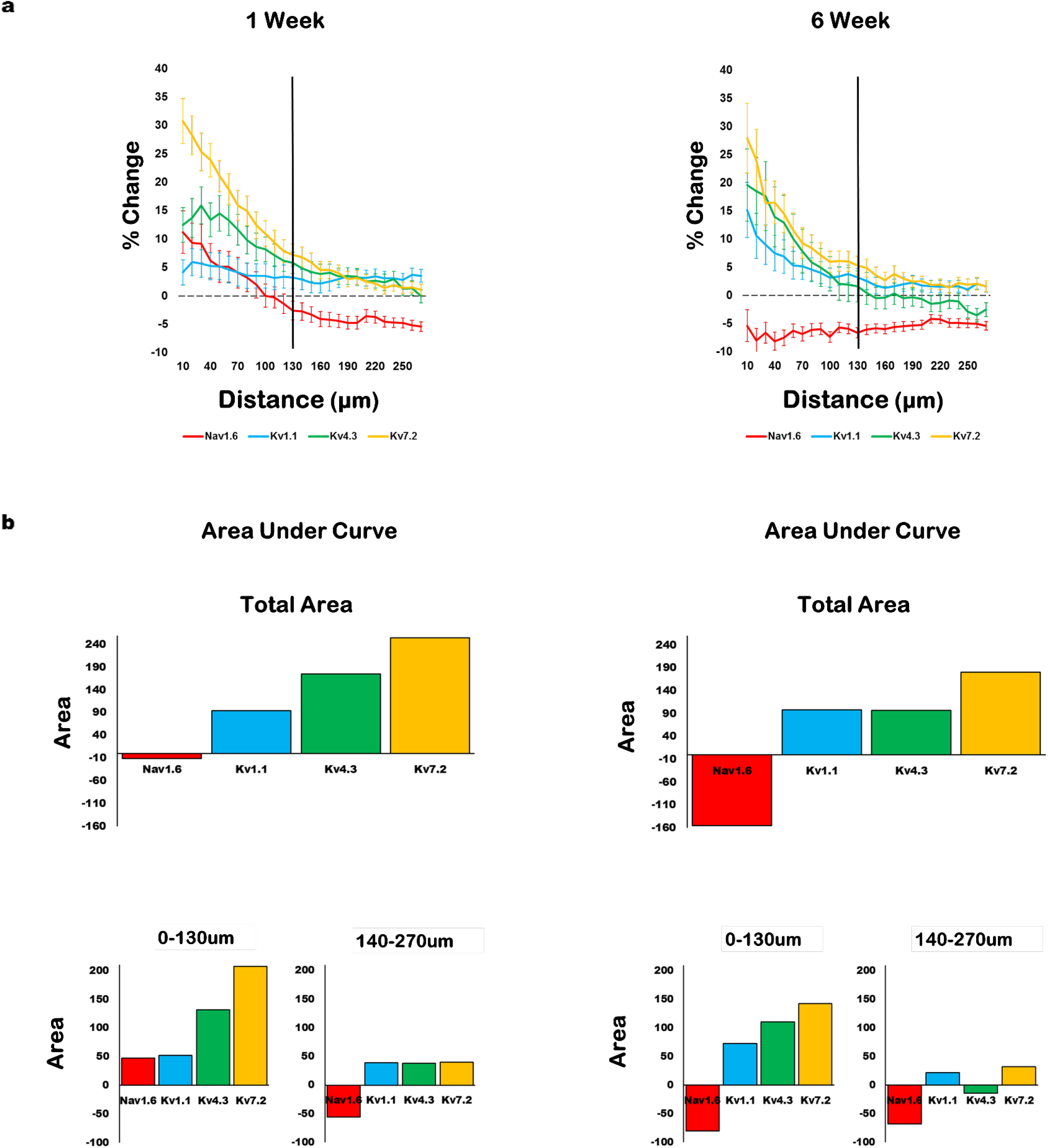
Temporal differences in expression: Percentage change in expression relative to 1 day values corroborates progressive reduction in sodium channel expression and heightened potassium channel expression over time. **A)** Averaged percentage change for 1 and 6 week expression values relative to 1 day expression values with standard error bars. **B)** Area under the curve calculated for unit region (0-130um) and LFP region (140-270um) for both 1 and 6 week expression curves, as well as total integrated area calculated for the combined 270um radius.

Since these total values did not appear to represent the interfacial differences observed (**Fig. 3A**), we further segmented the surveyed distance into two distinct regions to assess temporal shifts in expression levels within the estimated radius generating detectable single unit (0-130um)^30^ or LFP-only (140-270) activity (**Fig. 3A**). These distances were chosen based on the seminal work by Henze et. al, which determined the distances capable of producing sufficient amplitude for spike detection and clustering^30^. The results indicate variability in the time course of ion channel expression surrounding devices. At 1 week, the integrated area for the “unit” region revealed a relative increase in Nav1.6 (+47% integrated area), Kv1.1 (+52%), Kv4.3 (+132%), and Kv7.2 (+208%), while the integrated area for the “LFP” region revealed a relative decrease in Nav1.6 (−56%) and increase in Kv1.1 (+39%), Kv4.3 (+38%), and Kv7.2 (+40%). At 6 weeks, the “unit” region showed a decrease in Nav1.6 (−81%), and increase in Kv1.1 (+73%), Kv4.3 (+110%), and Kv7.2 (+143%). The integrated area for the “LFP” region had a decrease for both Nav1.6 (−68%) and Kv4.3 (−14%), and an increase in Kv1.1 (+22%) and Kv7.2 (+33%). Therefore, the relative shift in “unit” region Nav1.6 from elevation at 1 week to depression at 6 weeks, coupled with the sustained elevation in all Kv channels at both time points, indicates a temporal shift from hyper-to hypo-excitability within the recordable radius of the device relative to previous values.

### Alterations in ion channel expression accompany signal loss

Bi-weekly recordings taken across subjects demonstrated a progressive decline in single unit detection over 6 weeks (**Fig. 4A**). A relatively stable LFP amplitude experienced a decline at ~3 weeks that remained at a steady state over the remaining time course (**Fig. 4A**). To further investigate the relationship between ion channel expression and signal loss, we plotted ratios to explore relative interactions (**Fig. 4B**). The results revealed that Nav1.6/Kv7.2 expression ratio may be most predictive of unit loss, as the two metrics decrease in accordance with one another over 6 weeks (**Fig. 4**), whereas Nav1.6/Kv4.3 may be most predictive of LFP amplitude (**Fig. 4**). Nav1.6/Kv1.1, however, does not appear to correspond to either of the signal metrics. These results may provide insight into novel metrics for guiding device-tissue integration.

**Figure 4.**
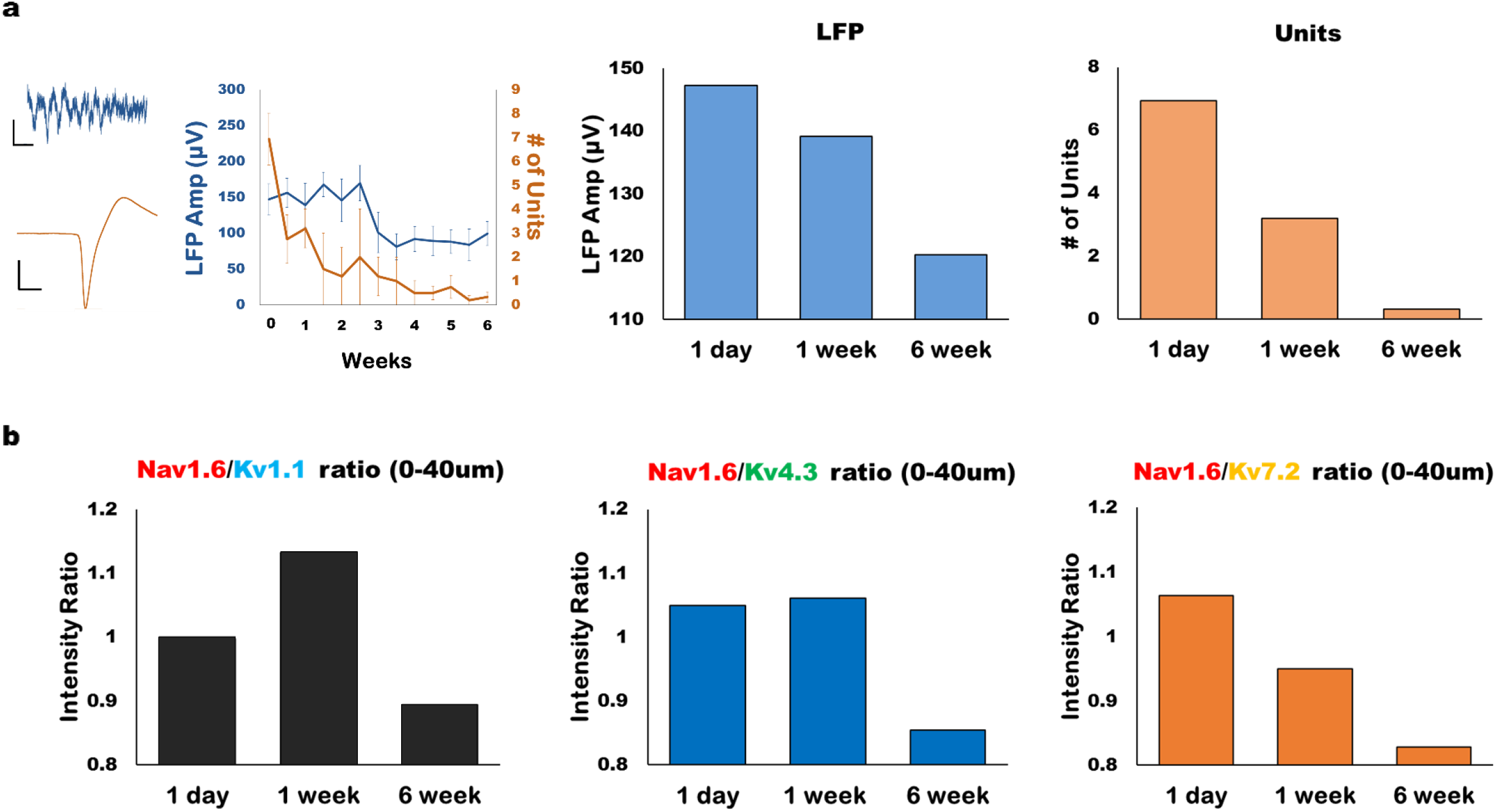
Alterations in ion channel expression accompany decline in unit detection. **A)** Example of putative unit and LFP snippet from microelectrode arrays, accompanied by the quantified data (# of units and LFP amplitude) obtained from bi-weekly recording sessions across subjects (with standard error bars). Average LFP amplitude and # of units plotted on bar graphs for each time point. **B)** Averaged data within 0-40um for intensity ratios are plotted. Nav1.6/Kv7.2 intensity ratio appears to coincide closest with unit detection over the 6 week time course, whereas Nav1.6/Kv4.3 ratio appears to best correspond to LFP amplitude over 6 weeks. In contrast, Nav1.6/Kv1.1 does not appear to correspond to either signal metric.

### Early observations suggest Kv7.2 expression modulates excitatory synaptic transporters

To explore whether ion channel expression may be a mechanism for shaping synaptic circuitry, we delivered siRNA in cultured rat cortical neurons to assess the consequences of Kv7.2 knockdown on excitatory synapses (to mimic transient reduction in Kv7.2 at 1 day, **Fig. 2A**). Neurons were transfected with either negative control siRNA (“scramble”) or siRNA against Kv7.2, and cells were harvested at either 3 or 7 days. RNA was collected to make cDNA, and primers for Kv7.2 (KCNQ2), VGLUT1, and PSD95 (post-synaptic density 95, an excitatory postsynaptic marker) were used to perform qPCR. Detection levels were normalized to scramble control levels for the respective primer. We observed elevations in VGLUT1 at 3 and 7 days when comparing Kv7.2 siRNA with negative control siRNA (**Fig. 5**). We observed a robust elevation in PSD95 at 3 days that was drastically reduced by 7 days (**Fig. 5**). These results suggest that Kv7.2, in accordance with previous reports^33^, regulates excitatory synaptic density (where previous reports demonstrated this relationship to VGLUT1 and PSD95 by pharmacological blockade of Kv7.2^33^). While preliminary, these results correspond with the in vivo results of VGLUT1 upregulation at 3 and 7 days (**Fig. 5**), using data from a previous report^41^. These results suggest that the transient reduction of Kv7.2 at 1 day *in vivo* (**Fig. 2A**) could contribute to the upregulation of VGLUT1 surrounding devices at 3 and 7 days (**Fig. 5A**)^41^.

**Figure 5.**
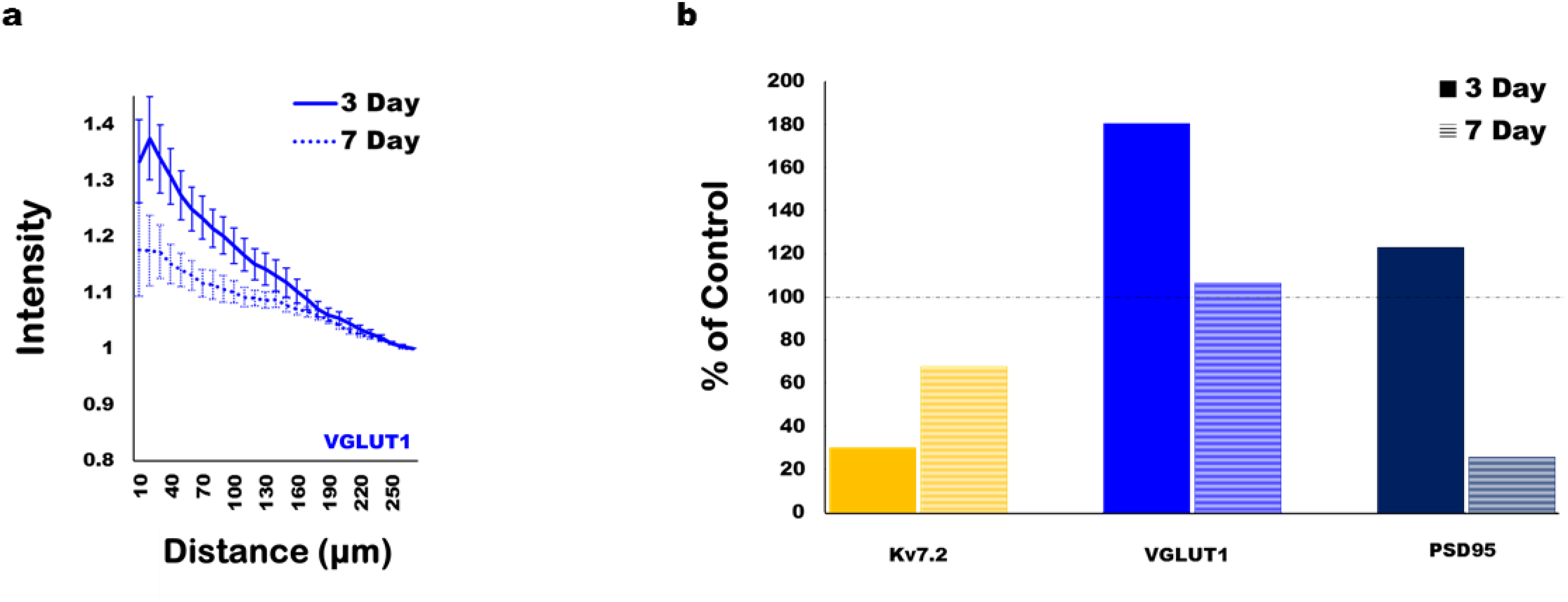
Preliminary observations suggest Kv7.2 knockdown impacts excitatory synapses in culture. **A)** *In vivo* results of vesicular glutamate transporter 1 (VGLUT1), using data from a previous report^41^, show an elevation in VGLUT1 at 3 and 7 days. **B)** *In vitro,* cortical neurons transfected with Kv7.2 siRNA show successful transient knockdown of Kv7.2, a similar trend in VGLUT elevation at 3 and 7 days compared to *in vivo* expression, and an impact on PSD95 in the form of a reduction at 7 days. Taken together, these data suggest that the transient downregulation of Kv7.2 at 1 day *in vivo* (**Fig. 2A**) may be a mechanism for the upregulation of VGLUT1 at 3 and 7 days *in vivo.* Two biological replicates were performed for the preliminary *in vitro* data.

## Discussion

Neuronal loss and glial encapsulation are traditionally used as metrics to assess the biocompatibility of devices for chronic neural interfacing^9,44–52^. However, recent work indicates that these conventional methods are insufficient to explain long-term signal quality^53^, where inter-day variability and progressive signal loss burden chronic recording arrays^3,5,6,54,55^. Well-characterized alterations in ion channels and synapses following cortical injury^10,12,22–25,35,56^ and inflammation^14–17,37,57^ suggest that similar alterations may accompany implanted devices. In fact, recent studies using non-functional microelectrode arrays have revealed changes in network *connectivity* (synaptic circuitry^41^) and *function* (calcium activity^58^) within the recordable radius of the device interface (~100um^30,59^), providing evidence of local circuit remodeling that may contribute to chronic signal instability. Here, we reveal changes in the fundamental components that underlie neuronal signaling (ion channels) within the recordable radius of the device interface^30,59^. The findings support our previously described trend from acute hyperexcitability to chronic hypoexcitability at the device interface^41^ and expand upon it by providing a potential link between ion channel and synaptic transporter expression (**Figs. 2 & 5**)^41^. Novel observations of ion channel expression surrounding devices revealed a progressive elevation in potassium and a reduction in sodium channel expression that temporally coincided with signal loss (**Figs. 2 & 4**). This work reveals insight into device-related mechanisms affecting the signal generation and firing properties (e.g., spike shape, firing rates, etc.) that underlie the characteristics of recorded signals.

The four ion channels were chosen based on their fundamental roles in regulating action potential generation^31^, firing patterns^32–34^, and synaptic efficacy^33^. Nav1.6, critical for action potential generation, has been implicated in electrophysiological abnormalities following axonal injury^35^, TBI^10,29^, and inflammation^17,36–38^. In addition, electrophysiological abnormalities following axonal injury can persist in both axotomized and neighboring intact neurons^29^. The authors attributed these electrophysiological abnormalities to changes in the expression of sodium channels^10,29,35^, where blocking sodium channel upregulation following TBI has been shown to improve outcomes by reducing excitability^13^. The authors additionally attributed abnormal activity to A-type potassium channels (with fast-activating/inactivating kinetics^60^), where reductions in channel expression has been shown to contribute to seizure susceptibility within initial days following TBI^12^ by increasing the excitability and firing rates of local neurons^12^. This is consistent with the transient downregulation of Kv4.3 observed at 1 day (**Fig. 2**), where the subsequent upregulation at 1 and 6 weeks may be a compensatory mechanism for counteracting hyperexcitability and epileptogensis. Combined, these data suggest that the reduction in Nav1.6 and upregulation of Kv4.3 at 6 weeks could inhibit action potential generation and dampen excitability/firing rates within the immediate vicinity of the implant. Kv1.1 upregulation due to CNS injury^39^ has been shown to likewise underlie axonal dysfunction in surviving axons, where increased K+ conductance was proposed to act as an axonal conduction block by shunting Na+ current^39,40^. This resulted in a reduction in the amplitude and area of compound action potentials for surviving axons at 6-8 weeks post-injury^39^. Therefore, the late upregulation of Kv1.1 observed at 6 weeks post-implantation may act as a shunt for preventing signal propagation within the recordable radius of the device-interface. Kv7.2 produces slowly activating and inactivating subthreshold M-currents, which are responsible for regulating excitability, responsiveness to synaptic inputs, and neuronal discharge frequency^61–63^. Kv7.2 channels located at pre- and post-synaptic terminals^62,63^ have been shown to be responsible for modulating neurotransmitter release, where M-current agonists prevent neurotransmitter release^64,65^. Therefore, an upregulation of Kv7.2 as observed at 1 and 6 weeks can reduce excitability, firing frequency and neurotransmission. Taken together, Nav1.6 reduction and Kv4.3 upregulation can limit the probability of action potential generation and dampen excitability/firing rate, Kv1.1 upregulation can provide excess shunt current to block downstream axonal conductance, and Kv7.2 upregulation can reduce responsiveness to synaptic inputs, inhibit repetitive firing and reduce neurotransmitter release at the synapse. Therefore, the reduced excitability and propagation/transmission of signals by ion channel alterations indicates a novel source for impaired signal detection by implanted recording arrays.

Signal loss over the 6 week time course was accompanied by a progressive elevation in potassium and reduction in sodium channel expression surrounding devices (**Figs. 2, 3 & 4**). At 1 day, the local reductions in Kv7.2 and Kv4.3 in the absence of effects on Nav1.6 or Kv1.1 may reflect a hyperexcitable state, which accompanied optimal unit detection (**Figs. 2 & 4**). The shift at 1 week to elevated Nav1.6, Kv4.3 and Kv7.2 coincided with a modest reduction in unit detection (**Figs. 2, 3 & 4**), which could be more heavily affected by the dual Kv4.3/Kv7.2 upregulations. The final shift at 6 weeks to a relative loss of Nav1.6 and gain in Kv1.1 indicates a more hypoexcitable state, which coincided with the poorest unit detection (**Fig. 4**). To further investigate this relationship, ion channel intensity ratios were plotted to compare with signal decline (**Fig. 4**). Nav1.6/Kv7.2 intensity ratio appears to temporally coincide best with unit detection. The decreased Nav1.6/Kv7.2 ratio indicates lower action potential probability from reduced sodium currents and increased sub-threshold K+ currents. Thus, the Nav1.6/Kv7.2 ratio could provide insight into neuronal excitability and firing rates that may contribute to unit loss. While the origin of the LFP was historically considered to largely emerge from postsynaptic potentials^66,67^, recent work indicates that it is instead mostly composed of non-synaptic currents^68^. Here, the Nav1.6/Kv4.3 ratio appears to correspond best with the LFP (**Fig. 4**), which coincides with modeling data showing that the LFP is dominated by active membrane currents rather than postsynaptic conductance changes^68^. Kv4.3 channels are critical for producing high-frequency activity (which is achieved by their fast inactivation recovery^60^). Enhanced activity from Kv4.3 upregulation could increase active membrane conductances, which could in turn attenuate LFP amplitude^68^. Moreover, the combined upregulation with Kv1.1 and Kv7.2 channels could also contribute to increased membrane leakiness that may underlie LFP attenuation by 6 weeks^68^ (**Fig. 4**). Finally, these results could potentially explain electrophysiological mechanisms that underlie inter-day variability of unit detection and amplitude^3,5,6,54,55^. For example, modeling data for ionic current contributions to extracellular action potentials demonstrate that conductance densities for heterogeneous subtypes of K+ currents largely underlie variability in recorded waveforms^69^. Thus, the fluctuations seen in Kv1.1, Kv4.3, and Kv7.2 across the 6 week time course could explain unit variability observed by chronic neural interfaces^3,5,6,54,55^. In addition, the fluctuations in Nav1.6 could likewise explain inter-day variability in amplitude^5,6,55,69^. Taken together, these results may provide novel metrics to assess the biocompatibility of devices for improved long-term function.

Our results must be interpreted relative to the well-known changes in cellular densities that are associated with chronically implanted electrodes, including neuronal loss and glial encapsulation^7,9,44–46,48,49^. However, density changes do not fully explain inadequate performance, day-to-day variability, and signal loss accompanied by ideal histology and device integrity^53^. Another important consideration is the potential for expression of ion channels to occur in non-neuronal cell types. Of the four ion channels assessed, the only channel expressed in non-neuronal cells (to the best of our knowledge) is Kv1.1, which is also expressed in microglia^70^. However, because the elevation in Kv1.1 did not occur until 6 weeks, this indicates that it is unlikely that microglia are the source of Kv1.1 expression, as a stark microglial layer forms around the device within initial hours and days^71^. The fact that Kv1.1 expression is stable at 1 day and 1 week across the observed 270um (**Figs. 2 & 3**), therefore, supports non-microglial labeling. In general, we observed subcellular expression patterns which were consistent with neuronal labeling. Kv1.1 appeared to be localized to axons and terminals as previously described^32,39^ (also validated with the vendor antibody^72^). Nav1.6 labeling appears consistent with somatic and axonal initial segment localization^35,42^, which aligns with previous reports using the same antibody^73,74^. Kv4.3 labeling is consistent with somatic localization in layer V pyramidal neurons^75^, and corresponds with validated labeling in hippocampal CA3 neurons using the vendor antibody^76^. Finally, Kv7.2 labeling appears to be expressed in axons and synaptic terminals^62,63^, where our specific antibody has been confirmed with heavy colocalization in the Nodes of Ranvier^77^. While these results will be further validated in future work, the staining appears to be consistent with neuronal localization.

Since Kv7.2 activity is known to regulate excitatory synaptic density (specifically VGLUT1 and PSD9 5^33^), we chose to explore the impact of Kv7.2 knockdown on excitatory synapses *in vitro.* Our preliminary results revealed upregulated VGLUT1 expression at both 3 and 7 days following Kv7.2 knockdown. These outcomes suggest that the reduced Kv7.2 expression observed at 1 day *in vivo* may be a mechanism for upregulating VGLUT1 expression at 3 and 7 days *in vivo* (**Figs. 2 & 5**) as previously reported^41^. The subsequent reduction in PSD95 at 7 days may be initiated by excitotoxicity at earlier time points. Since glutamate release scales with VGLUT1 expression^78^, excessive glutamate release (coupled with hyperexcitability) could explain the loss of PSD95 (where dramatic decreases in PSD95 have been shown in excitotoxic models^79^). Therefore, the observed trend toward hypoexcitability (**Fig. 2**) could be a reparative effort to promote neuroprotection and prevent further excitotoxicity. While acute alterations in potassium channel expression may be responsible for the shift in synaptic circuitry *in vivo*, the underpinnings responsible for the shift in ion channel expression will need to be identified in future work. Sources may include reactive signaling cascades initiated by insertion (such as the release of inflammatory cytokines that alter ion channel expression and function)^9,17^, and strategies to modify inflammatory mechanisms have improved long-term recording quality in previous reports^80,81^. This work may provide new insight into mechanisms of tissue reactivity surrounding devices that may contribute to signal loss.

Next-generation device designs are emerging to tune the tissue response to mitigate gliosis and neuronal loss^44–49^ in an effort to develop recording arrays with improved long-term function. However, ideal histology and device integrity based on these traditional methods have still not guaranteed adequate recording quality^53^, suggesting that the principles guiding the design of improved devices may require further consideration. By assessing the fundamental components that underlie neuronal signaling (ion channels and synaptic circuitry), the innovative methods described herein may provide a more reliable indication of recorded signal quality based on their inherent contributions to neuronal signaling events. We have provided four (4) fundamental ion channels that appear to be especially informative of recording quality based on their corresponding relationships (**Figs. 2, 3 & 4**). Specifically, the number of units detected over 6 weeks appears to correspond best with the Nav1.6/Kv7.2 ratio (**Fig. 4**), and LFP amplitude appears to correspond most closely with the Nav1.6/Kv4.3 ratio (**Fig. 4**). This technique can be implemented to not only guide next-generation device designs (e.g., architecture, size, flexibility, surface chemistry/topography^9,44–47^), but also intervention strategies (e.g., coatings, microfluidic delivery, etc.^82–85^) aimed at improving long-term recording quality.

## Methods

### Surgery

Adult male Sprague-Dawley rats (SAS, 250-400g, Charles River, Wilmington, MA) were bilaterally implanted in the primary motor cortex with 16-channel single-shank microelectrode arrays (A1×16-3mm, 703um^2^ site sizes, NeuroNexus, Ann Arbor, MI) using a surgical procedure similar to that previously described^41^. Briefly, animals were anesthetized and maintained at ~2.0% isoflurane throughout surgery, whereby a 2×2mm craniotomy was performed over the primary motor cortex (+3.0mm AP, 2.5 ML), the dura was resected, and a single-shank probe was stereotaxically inserted 2mm from the cortical surface. Dental acrylic was used to secure the bilateral implants, where a bone screw was placed posterior of each device to anchor the headcap. Bupivacaine and Neosporin were topically applied around the wound to minimize discomfort and risk of infection, and meloxicam was administered for pain management. All surgical procedures were approved by the Michigan State University Animal Care and Use Committee.

### Extracellular electrophysiology

Bi-weekly recording sessions were performed with isoflurane (~1-1.5%) using TDT software (Tucker Davis Technologies, TDT, Alachua, FL) by connecting a ZIF-clip headstage to a Z25 pre-amplifier (TDT) and PZ2 amplifier (TDT), to obtain 5 minute recording blocks per device per recording session. Low-pass filter for local field potential (LFP, 300Hz) and high bandpass filter for unit activity (500Hz-5KHz), yielded recording blocks that were then analyzed using a previously reported MATLAB script^8,46^ to determine the LFP amplitude and number of units. Single units were detected based on threshold crossings (3.5 standard deviations from noise floor), where principal component analysis and fuzzy c-means clustering were then used to isolate putative units (in combination with visual inspection of mean waveforms).

### Histology

Animals were deeply anesthetized using sodium pentobarbital at predetermined time points (24hrs, 1wk, 6wks) and transcardially perfused with PBS followed by 4% PFA. Explanted brains were postfixed overnight in 4% PFA at 4°C, and then sucrose protected for cryoembedding. Immunohistochemistry was performed according to previously reported methods^41^, where 20μm-thick horizontal cryosections from depths estimated in layer V of primary motor cortex were hydrated in PBS, blocked in 10% normal goat serum (NGS) in PBS and subsequently incubated in primary antibodies overnight at 4°C. The sections were rinsed the following day with PBS, incubated with secondary antibodies, and coverslipped with ProLong Gold antifade reagent (Molecular Probes by Life Technologies, Carlsbad, CA). Antibodies were diluted in carrier solution consisting of 5% NGS and 0.3% Triton X-100 in PBS. Primary antibodies included rabbit anti-Nav1.6, −Kv1.1, −Kv4.3, and −Kv7.2 (1:200, Alomone Labs, Jerusalem, Israel). Secondary antibodies included goat anti-rabbit IgG (H+L) alexa fluor 594 conjugate (1:200, Thermo Fisher Scientific, Waltham, MA). An Olympus Fluoview 1000 inverted confocal microscope was used to image samples with a 20x PlanFluor dry objective (0.5NA), where settings were optimized for individual images as previously described^86^. Images were then analyzed with a previously reported MATLAB script^41^ adapted from Kozai et. al^86^. Briefly, 10um concentric bins were generated to radiate concentrically from a user-drawn injury outline (a total of 27 bins spanning a 270um radius), where the pixel intensity was averaged within each bin. In this way, image intensity was analyzed as a function of distance to quantify interfacial patterns of protein expression over distance and time. Area under the curve was calculated using the *trapz* function in MATLAB to perform discrete integration on the averaged intensity data points.

### Cell culture and transfection

Rat primary cortical neurons (E18, Life Technologies, Carlsbad, CA) were cultured in neurobasal medium (1mL B27, 125 uL GlutaMax in 50mL Neurobasal Media) for one week prior to transfection. For transient transfections, siRNA (Kv7.2 or negative control stealth, Life Technologies, Carlsbad, CA) was mixed with Optimem and Lipofectamine RNAiMax (according to manufacturer’s instructions) and incubated overnight, followed by a complete exchange with fresh neurobasal media. Cells were harvested after 3 or 7 days post-transfection (RNEasy mini kit, Qiagen), whereby cDNA was made and amplified via qPCR with primers for GAPDH, KCNQ2 (Kv7.2), VGLUT1, and PSD95. All primer levels were normalized to GAPDH levels, and then normalized to the scramble siRNA control levels for each primer.

### Statistical analysis

A linear mixed effects model was performed with SPSS (IBM, Chicago, IL) and incorporated both distance and temporal effects. Results were assessed using a Fischer’s Least Significance Difference test and defined as significant at *p<0.05 and ***p<0.001. For each ion channel, there was an average of 7 devices and 21 tissue sections analyzed per time point. At 1 day, there was an average of 5 devices and 12 tissue sections analyzed per ion channel stain; at 1 week, an average of 9 devices and 30 tissue sections; and at 6 weeks, an average of 7 devices and 21 tissue sections.

## Acknowledgments

This work was supported by NIH grant 1R21NS094900 (NINDS), and the Departments of Biomedical Engineering and Electrical and Computer Engineering at Michigan State University. Thanks to Steven Suhr of Biomilab, LLC for *in vitro* knockdown training, Melinda Frame from Center for Advanced Microscopy for confocal training, Stefanos Palestis for assistance with signal processing, Matthew Drazin for assistance with cyrosectioning, and TK Kozai and Bailey Winter for the intensity profiling MATLAB script. Thanks to TK Kozai and Ali Mohebi for valuable feedback.

